# Nutrient Responsive O-GlcNAcylation Dynamically Modulates Galectin 3 Secretion

**DOI:** 10.1101/2021.04.05.438483

**Authors:** Mohit P. Mathew, Julie G. Donaldson, John A. Hanover

**Author notes:** Corresponding Author: John A. Hanover, Bldg 8 room B127, NIH 9000 Rockville Pike, Bethesda Md 21895, E mail.

## Abstract

Endomembrane glycosylation and cytoplasmic O-GlcNAcylation each play essential roles in nutrient sensing, and in fact, characteristic changes in glycan patterns have been described in disease states such as diabetes and cancer. These changes in glycosylation have important functional roles and can drive disease progression. However, little is known about the molecular mechanisms underlying how these signals are integrated and transduced into biological effects. Galectins are proteins that bind glycans that are secreted by a poorly characterized non-classical secretory mechanism. Once outside the cell, galectins bind to terminal galactose residues of cell surface glycans and modulate numerous extracellular functions like clathrin independent endocytosis (CIE). Originating in the cytoplasm, galectins are predicted substrates for O-GlcNAc addition and removal. This study shows that galectin 3 is O-GlcNAcylated, and that changes in O-GlcNAc cycling alters its secretion. Moreover, we determined that there is a significant difference in O-GlcNAcylation status between cytoplasmic and secreted galectin 3. We observed dramatic alterations in galectin 3 secretion in response to nutrient conditions and that these changes were dependent on dynamic O-GlcNAcylation. Finally, we showed that alterations in galectin 3 secretion via disrupted O-GlcNAcylation drove changes in CIE. These results indicate that dynamic O-GlcNAcylation of galectin 3 plays a role in modulating its secretion and can tune its function of transducing nutrient sensing information coded in cell surface glycosylation into biological effects.

## INTRODUCTION

Nutrient sensing is an essential function of glycosylation(1–4). This role is especially important in the context of disease where metabolic imbalances can be detected via alterations in patterns of glycosylation(5–10). Glycan synthesis is non-template directed therefore glycan structures are dependent on the availability of their carbohydrate components (i.e. sugars). As a result, glycosylation allows cells to ‘sense’ the availability and abundance of nutrients (in particular various sugars) and present and display this information in the form of altered glycan patterns. However, it is not well understood how this information is transduced into functional responses. Galectin 3 is a unique protein that provides a link between sensing changes in glycosylation and modulating cellular functions. Galectin 3 recognizes and binds to carbohydrates and hence can detect changes in glycan patterns(11–14). It has also been shown to mediate several cellular functions including clathrin independent endocytosis (CIE)(15–25). The position of galectin 3 at the interface of glycan pattern detection and modulating cellular function could suggest an important role for galectin 3 as a functional transducer in nutrient sensing.

Galectin 3 is a lectin that is synthesized in the cytoplasm and secreted by ‘non-canonical’ secretion(12,26–28). Galectin 3 binds to terminal galactose residues and is therefore able to detect changes in cell surface glycosylation patterns (which is one way in which cells detect and present nutrient information), galectin 3 is unique among the galectin family of proteins due to its ability to multimerize into pentameric complexes.(12–14). It has also been shown to drive a number of cellular functions including CIE and cell spreading(15–17,21–23,29,30). These observations uniquely position galectin 3 as a link between detecting nutrient sensing information and functional outcomes. A key element of this function of galectin 3 is that it is an extracellular function and hence requires secretion. Currently, there is little known about how this protein gets across the plasma membrane(26, 31). While export via exosomes has been proposed as a means of its secretion(32), a recent genome-wide CRISPR screen for galectin 3 secretory machinery demonstrated that secreted galectin 3 was primarily free and not exosome bound, suggesting that exosomal export was not the primary secretory pathway(33).

Since galectin 3 is synthesized in the cytoplasm it might be *O-*GlcNAcylated. *O-*GlcNAcylation is a ubiquitous post translational modification of cytoplasmic proteins. It involves the addition of a single N-acetylglucosamine sugar onto a serine or threonine residue of a protein. There are 2 enzymes involved in this modification, *O-*GlcNAc transferase (OGT) which adds a GlcNAc residue to a protein and *O-*GlcNAcase (OGA) which removes the *O-*GlcNAc modification (34–37). This modification has been observed on over 1000 proteins and has been shown to dynamically modulate protein signaling and stability in ways that are analogous to phosphorylation (34–37). Importantly, the sugar donor for *O-*GlcNAcylation (UDP-GlcNAc) is not only dependent on glucose metabolism but is also be influenced by cellular amino acid, fatty acid and nucleotide levels. Thus, *O-*GlcNAcylation plays an important role in nutrient sensing(1,2,35,36).

This study was designed to determine if galectin-3 is O-GlcNAcylated and to examine the effects of the O-GlcNAc modification on galectin-3 secretion and function. Our data suggest that galectin-3 can serve as a nexus between nutrient sensing information represented by cell surface glycosylation and intracellular O-GlcNAcylation, integrating information from both these processes and transducing it into functional biological outcomes (**Figure 1**).

**Figure 1:**
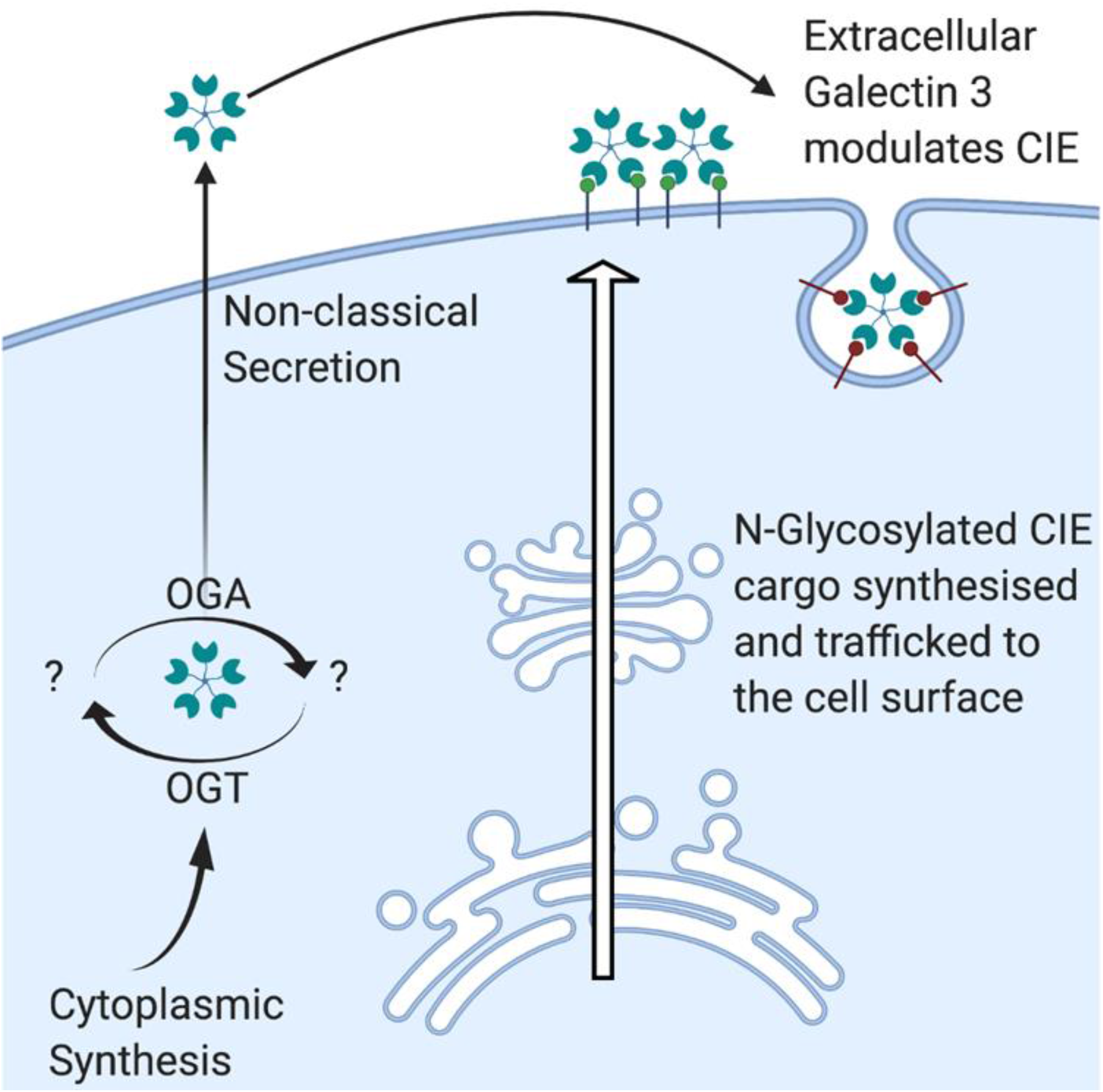
Schematic depiction of the role of galectin 3 in nutrient sensing. Galectin 3 is synthesized in the cytoplasm and then secreted by non-canonical mechanisms. Once outside the cell is binds to terminal galactose residues and can modulate a number of important cellular functions like clathrin independent endocytosis, cell spreading, etc. N-glycosylation is a non-template directed process and as such the glycan patterns that are produced are dependent on the availability of the enzymes and the substrates, as a result changes in nutrient environment are presented as altered glycan patterns. These glycan patterns are detected and bound to by galectins, which can then transduce these changes in pattern into changes in cellular behavior via the processes these galectins modulate. The cytoplasmic origin of galectins allows the possibility that these proteins are O-GlcNAcylated. O-GlcNAcylation also plays a role in nutrient sensing and is known to dynamically modulate the behavior of numerous proteins. This raises the potential for galectin 3 to integrate nutrient sensing information from both O-GlcNAcylation and N-Glycosylation and then transduce this information into biological effects.

## RESULTS

### Galectin 3 is O-GlcNAcylated

Based on the cytoplasmic synthesis of galectin 3 and its primary sequence, we hypothesized that it could be post translationally modified by O-GlcNAcylation. To test this hypothesis, we immunoprecipitated galectin 3 from Hela cells with and without GlcNAc treatment and western blotted for O-GlcNAcylated proteins using the RL-2 antibody (**Figure 2A**). We found an RL-2-positive band overlapping the Galectin 3 band suggesting that galectin 3 is O-GlcNAcylated. Surprisingly we also found a number of other O-GlcNAcylated proteins that co-immunoprecipitated out with galectin 3. In order to identify these co-immunoprecipitating proteins and to attempt to identify specific sites of O-GlcNAcylation on Galectin 3 we immunoprecipitated Galectin 3 and ran electron transfer dissociation (ETD) mass spectrometry (**Figure 2B**). While the mass spectrometry analysis wasn’t able to identify specific sites of O-GlcNAcylation on Galectin 3, it did identify 12 interacting proteins that were O-GlcNAcylated (**Table 1**). These interacting proteins included Sec 24C, which is a member of the COPII complex that has preciously been shown to be functionally regulated by O-GlcNAcylation(38).

**Figure 2:**
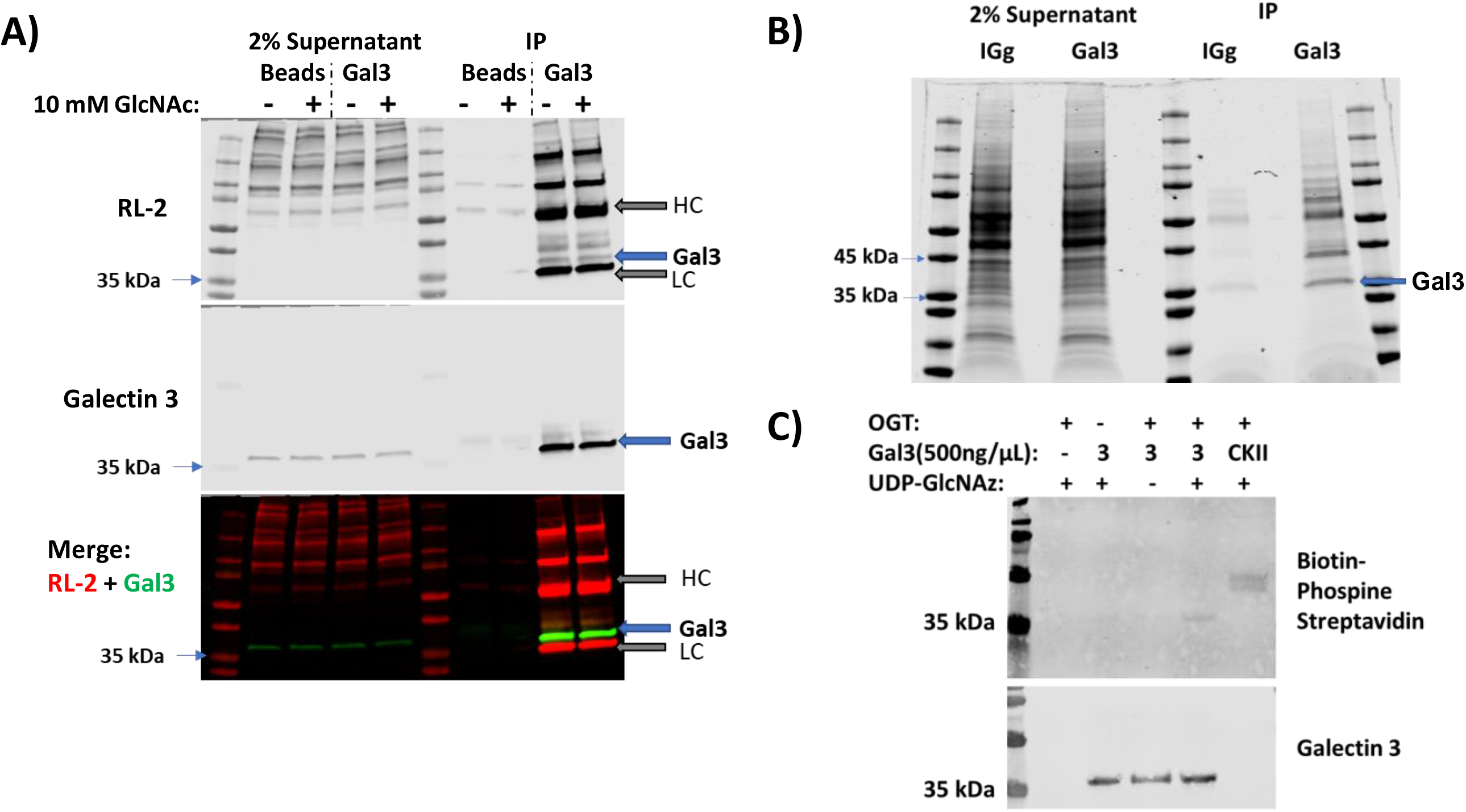
Galectin 3 is O-GlcNAcylated. Immunoprecipitation of galectin 3 from cells with or without 10mM GlcNAc treatment followed by western blotting shows an overlapping band of anti-galectin 3 and RL-2 binding indicating that galectin 3 is O-GlcNAcylated (**A**). A number of other RL-2 labelled bands were also pulled down suggesting that galectin 3 is also interacting with a number of O-GlcNAcylated proteins. Immunoprecipitated galectin 3 and interacting protein bands (**B**) were analyzed by mass spectrometry. An *in vitro* O-GlcNAcylation assay using recombinant galectin 3, recombinant OGT and UDP-GlcNAz confirms that galectin 3 can be O-GlcNAcylated (**C**). At least three independent experiments were carried out with representative blots shown.

**Table 1:**
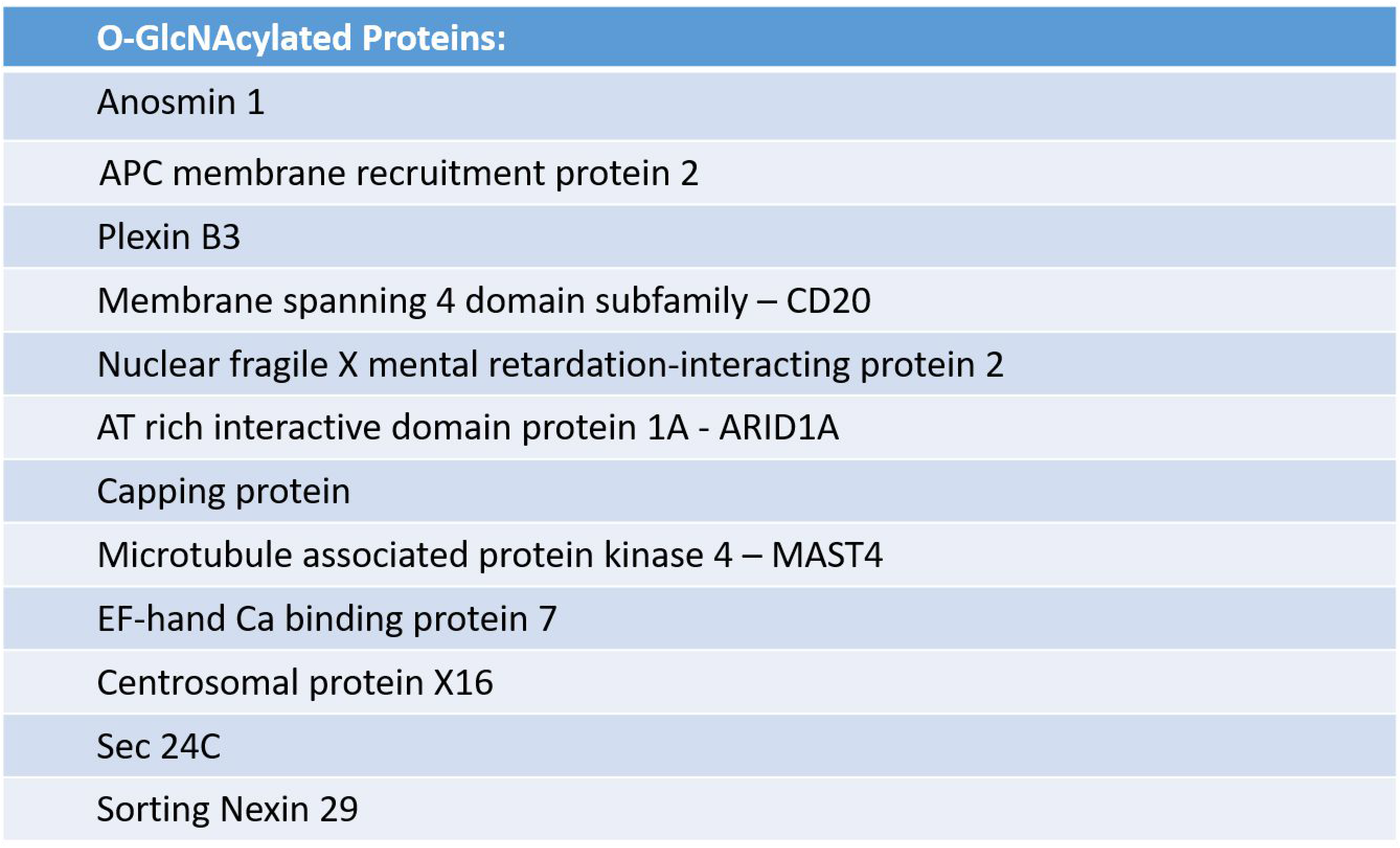
A list of 12 proteins were identified as O-GlcNAcylated by mass spectrometry.

In order to further validate the O-GlcNAcylation of galectin 3 we performed an *in vitro* O-GlcNAcylation assay wherein we incubated recombinant galectin 3 with UDP-GlcNAz and recombinant OGT, we then labelled the azide groups with biotin phosphine ran the product out on a western blot and blotted with streptavidin (**Figure 2C**). The recombinant galectin 3 was O-GlcNAcylated *in vitro* similar to the positive control. WGA pull down from Hela cells followed by competitive release using GlcNAc also pulled down galectin 3 further strengthening the case that galectin 3 is O-GlcNAcylated (**supplementary figure 1**).

### Cellular and secreted pools of galectin 3 are differentially O-GlcNAcylated

Next we overexpressed Galectin 3-GFP in Hela cells. After 3 days of transfection, galectin 3-GFP was pulled down from both the cellular and cytoplasmic pools of proteins using the GFPtrap system and was then western blotted (**Figure 3A**). RL-2 staining showed a clear overlapping band with the galectin 3-GFP band in the cellular pull down and input lanes confirming that galectin 3 is O-GlcNAcylated. While the inputs in the secreted pool were too dilute to be detected by western blotting, we were able to detect the pulled down band of secreted galectin 3 GFP. Interestingly, there was very little RL-2 staining corresponding to the secreted galectin-3-GFP pull down bands. Relative quantification normalizing RL-2 staining to the level of galectin 3 confirmed that there was a significant decrease in O-GlcNAcylation level in the secreted pool of galectin 3 (**Figure 3B**). These results indicate that the O-GlcNAcylation of galectin 3 may play a role in its secretion and that the removal of O-glcNAcylation is an important step before galectin 3 can be secreted.

**Figure 3:**
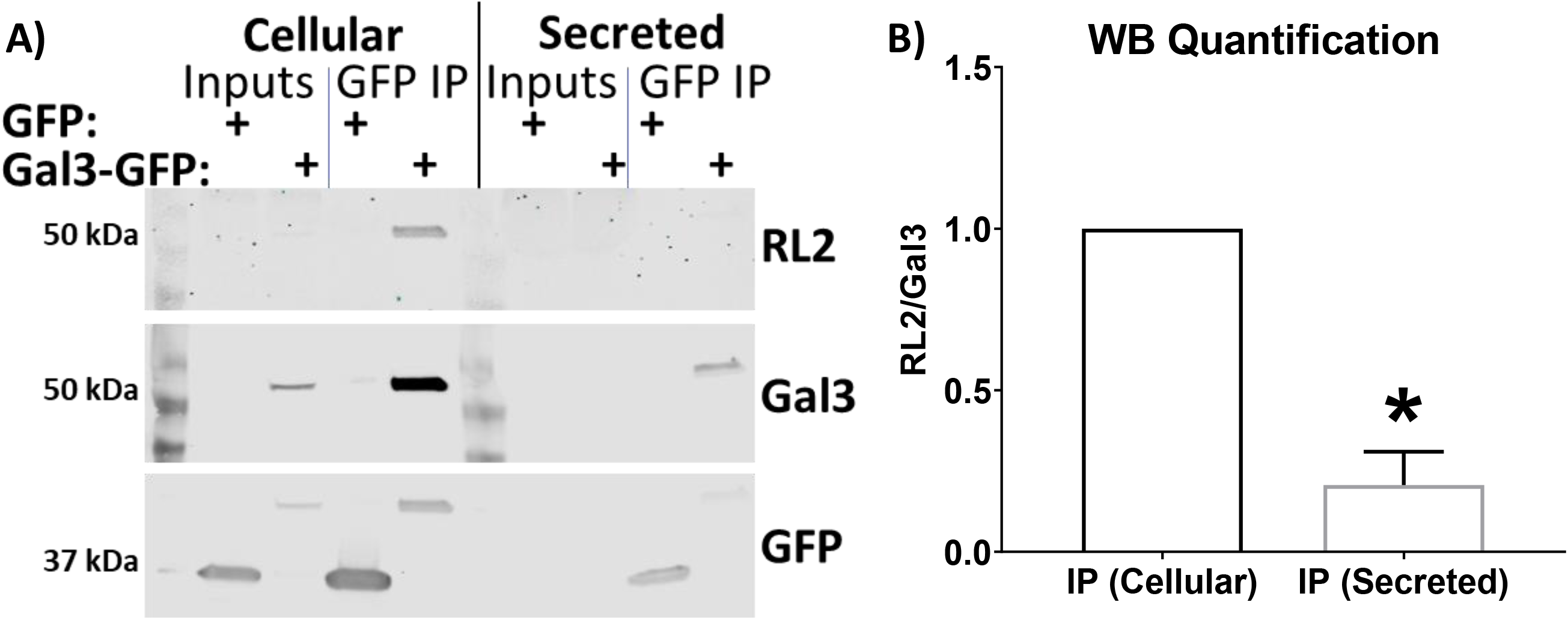
Secreted Galectin 3 is preferentially deglycosylated. Galectin 3-GFP was overexpressed in hela cells, galectin 3 from the cells and the conditioned media was then immunoprecipitated using the GFP-TRAP system (**A**). Quantifying and normalizing the RL-2 band to the galectin 3 band confirms that secreted galectin 3 is significantly less O-GlcNAcylated (**B**). At least three independent experiments were carried out with representative blots shown and data expressed as mean ± S.D. (error bars). *, p < 0.05.

### OGA KO MEFs secreted significantly less galectin 3

To study the effect of O-GlcNAcylation on galectin 3 secretion we turned to a genetic model of mouse embryonic fibroblasts from OGA knockout mice. Knockout of OGA prevents the removal of O-glcNAc residues and leads to an increase in O-GlcNAcylation levels. Using an ELISA assay to measure the amount of galectin 3 secreted per cell over the course of a 3 day incubation we found that in both immortalized and primary MEFS there was a significant decrease in galectin 3 secretion in OGA KO MEFS (**Figure 4**). This inhibition in secretion was observed in MEFs from both male and female mice although there did appear to be a slightly more dramatic effect in the male MEF cell lines.

### Disruption of O-GlcNAc cycling in Hela cells using SiRNA Knockdowns alters galectin 3 secretion

We then used siRNA to Knock down either OGT or OGA in Hela cells. Both OGA and OGT knockdowns resulted in a knockdown of OGA and an increase in O-GlcNAcylation levels (possibly due to the O-GlcNAcylation dependent regulation of these proteins themselves) (**Figure 5A**). Galectin 3 secretion over a 3-day incubation was measured using an ELISA assay. In both these knockdowns we observed a significant decrease of galectin 3 secretion consistent with what we had observed in the OGA KO MEFS (**Figure 5B**).

**Figure 4:**
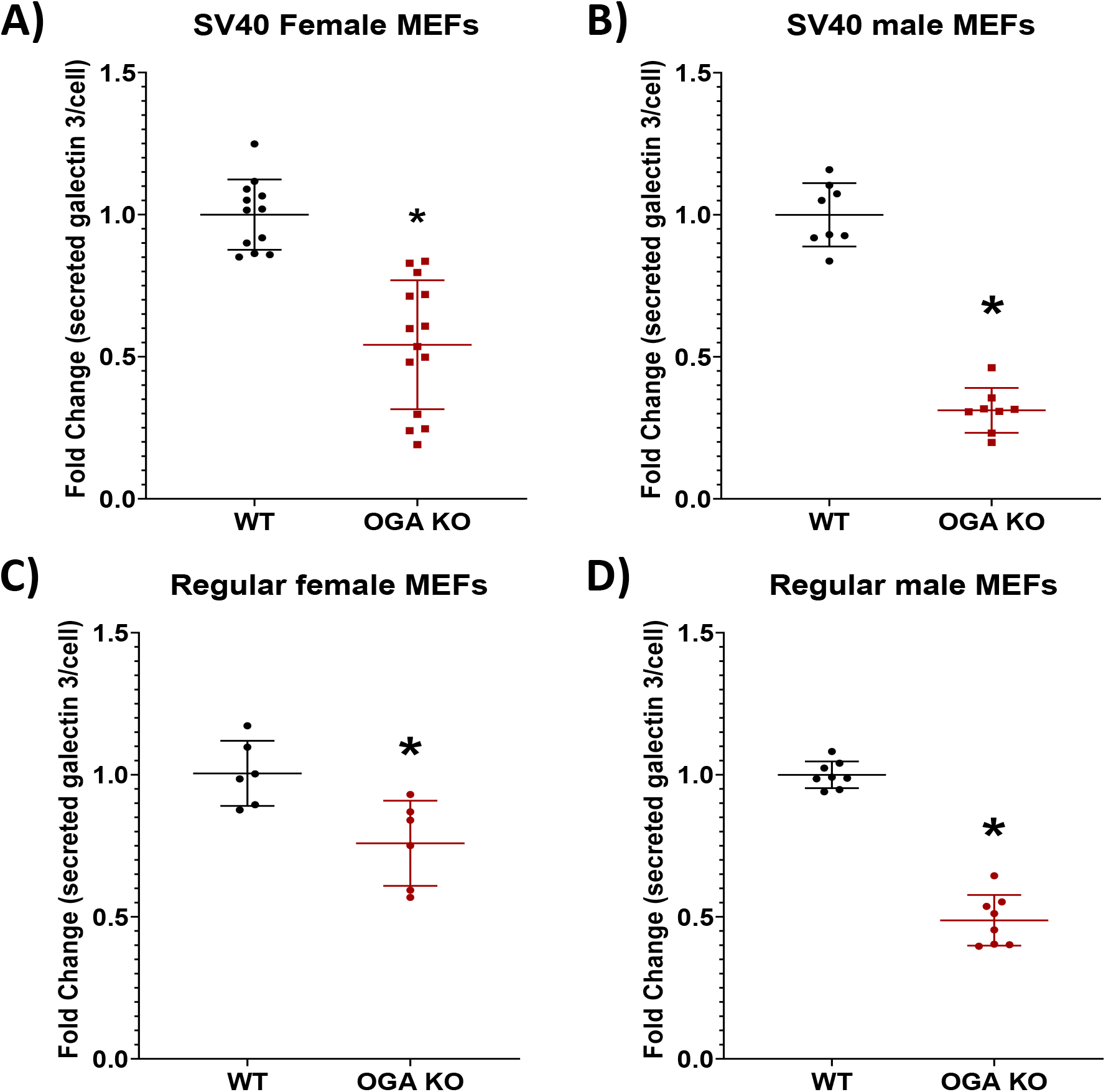
Mouse embryonic Fibroblasts from OGA Knock Out mice secrete significantly less galectin 3. Galectin 3 secretion was measured using an ELISA assay. Knock Out MEFS secreted less galectin 3 than wild type MEFs, this was observed in MEFS obtained from female mice (**A and C**), male mice (**B and D**). This secretion defect was also observed in both immortalized (**A and B**) and non-immortalized MEFS (**C and D**). At least three independent experiments were carried out with data expressed as mean ± S.D. (error bars). *, p < 0.05.

**Figure 5:**
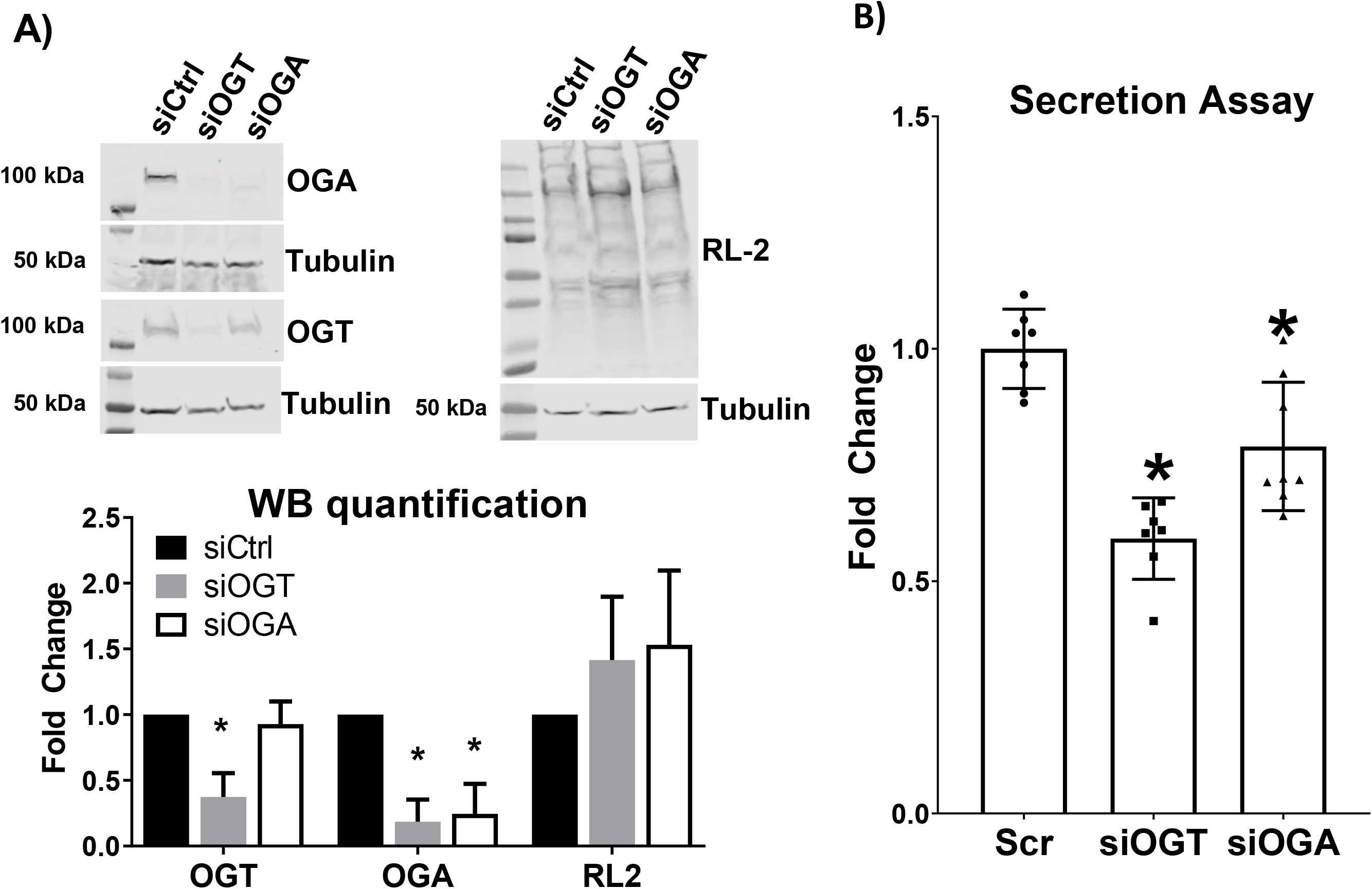
Disruption of O-GlcNAc cycling by SiRNA knockdown of OGT and OGA alters galectin 3 secretion. Western blotting shows that knocking down OGT or OGA using siRNA transfection leads to decreased OGA levels (OGA knockdown) or a decrease in both OGA and OGT levels (OGT knock down) in both cases there was an increase in O-GlcNAcylation observed (RL-2 binding) (**A**). Increased O-GlcNAcylation by knocking down either OGA or OGT leads to a decrease in galectin 3 secretion measured by an ELISA assay (**B**). At least three independent experiments were carried out with representative blots shown and data expressed as mean ± S.D. (error bars). *, p < 0.05.

### Chemical inhibition of O-GlcNAc cycling in Hela cells also alters galectin 3 secretion

Next, we used chemical inhibitors to disrupt O-GlcNAc cycling. Thiamet G inhibits OGA and led to increased levels of O-GlcNAcylation and OSM1 inhibits OGT and led to decreased levels of O-GlcNAcylation (**Figure 6A**). Galectin 3 secretion over the course of 2 days wasn’t affected by Thiamet G and increased significantly with OSM1 treatment (supplementary figure 2). Galectin 3 secretion over the course of 3-days showed a slight but statistically significant increase with Thiamet G treatment and a dramatic over 2-fold increase in secretion with OSM1 treatment (**Figure 6B**).

**Figure 6:**
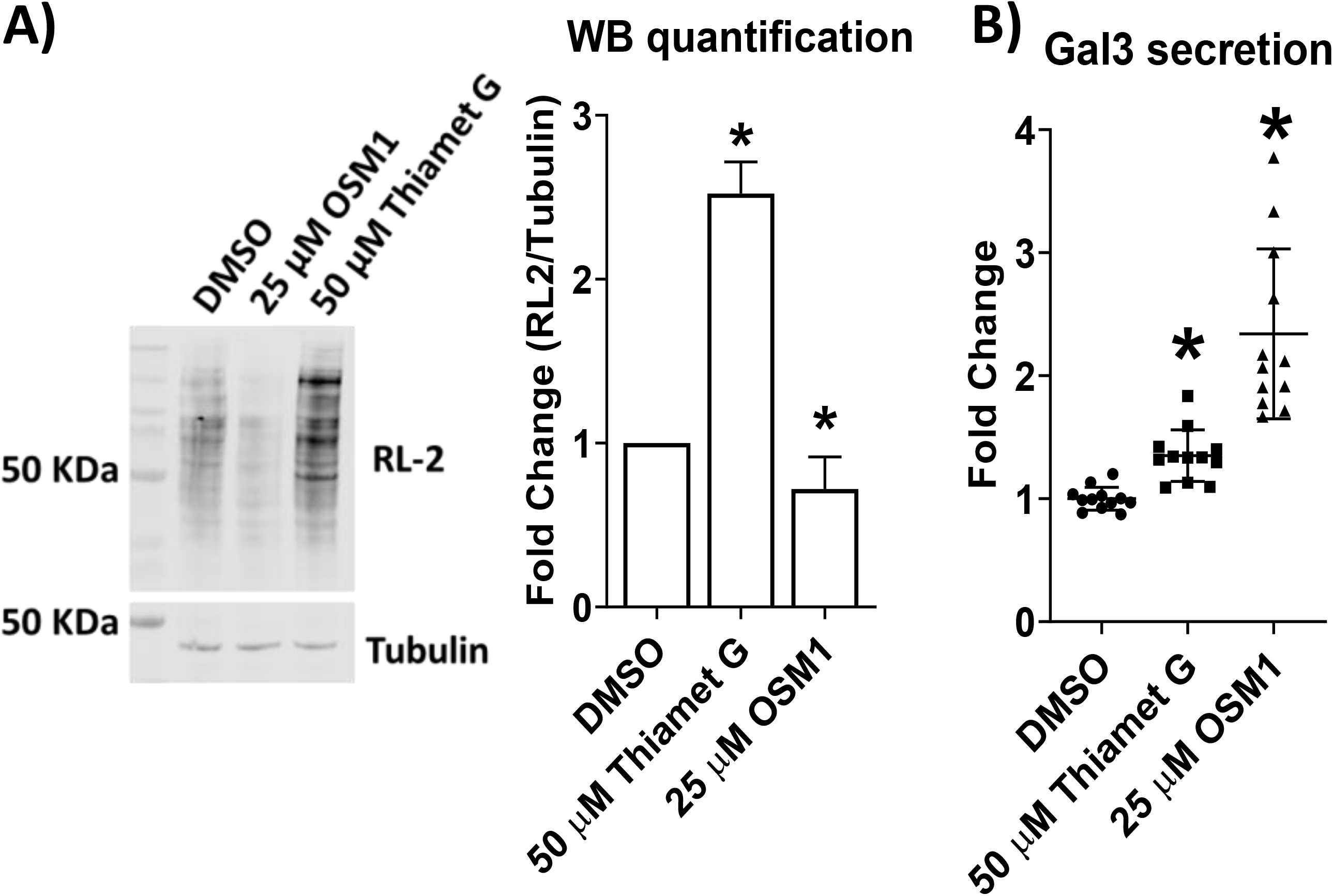
Chemical inhibition of OGT and OGA also affects galectin 3 secretion. Western blotting shows that inhibiting OGT and OGA using OSM-1 and Thiamet G over a 3-day incubation respectively resulted in a decrease in O-GlcNAcylation observed (RL-2 binding) for OSM-1 treatment and an increase in O-GlcNAcylation observed (RL-2 binding) for Thiamet G treatment (**A**). An ELISA based secretion assay shows that OSM-1 treatment led to a significant increase in galectin 3 secretion whereas Thiamet G treatment resulted in a small increase in secretion (**B**). At least three independent experiments were carried out with representative blots shown and data expressed as mean ± S.D. (error bars). *, p < 0.05.

These results indicate that Knockdown of OGA inhibits the secretion of galectin 3. The combination of data from OGT and OGA knockdown and OSM1 inhibition suggests that increased O-GlcNAcylation inhibits galectin 3 secretion whereas decreased O-GlcNAcylation stimulates galectin 3 secretion potentially by a sequestration-release model. A possible mechanism is that O-GlcNAcylation of galectin 3 signals its sequestration inside the cell and its deglycosylation is required to release it from the cell this is also consistent with the galectin 3-GFP pull down assay that showed a clear decrease in O-GlcNAcylated galectin 3 in the secreted form. The thiamet G data however suggests that this regulation is not entirely straightforward and may be a combination of direct and indirect effects.

### Direct disruption of galectin 3 O-GlcNAc cycling using mutant galectin 3-GFP alters its secretion

Next, we analyzed the protein sequence of galectin 3 for predicted sites of O-GlcNAcylation using the YinOYang server (**Figure 7A**). There were 6 sites of O-GlcNAcylation predicted with a high degree of confidence. Five of these predicted sites clustered in the multimerization region of galectin 3 while the last one (Ser 243) is adjacent to the nuclear export region.

**Figure 7:**
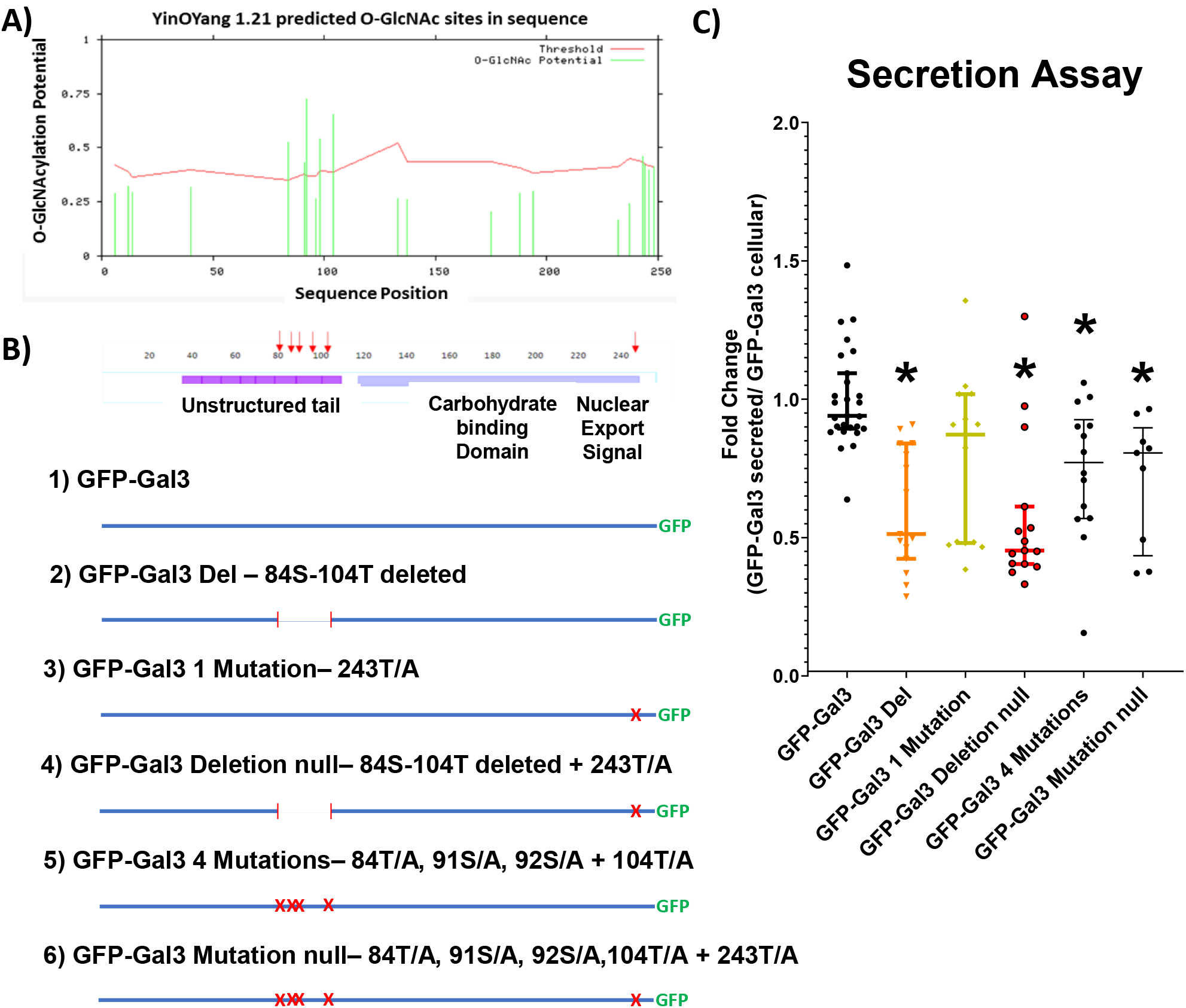
O-GlcNAcylation site mutants show direct defects in secretion. Sequence analysis of galectin 3 using the YinOYang software identified 5 predicted sites of O-GlcNAcylation (**A**). (**B**) Schematic representation of site mutants generated. Fluorescence based secretion assay confirms that mutant constructs with either a deletion or mutations of all the sites in the unstructured tail region result in significant secretion defects (**C**). At least three independent experiments were carried out and data expressed as mean ± S.D. (error bars). *, p < 0.05.

In order to better understand the direct role of O-GlcNAcylation on galectin 3, i.e. how changes in O-GlcNAcylation of galectin 3 itself affects its secretion, we prepared a set of galectin 3-GFP mutants with various combinations of predicted O-GlcNAcylation sites mutated (**Figure 7B** and **supplementary figure s3A**). These mutants were then expressed in Hela cells and after a 3 day incubation the fraction of galectin 3-GFP secreted was measured using a fluorescent plate reader assay (**Figure 7C**). All the mutants in which all 5 sites in the multimerization region are deleted or site mutated were secreted at a significantly lower level than control galectin 3-GFP. The single mutation near the nuclear transport region did not appear to alter secretion on its own. This would suggest that O-GlcNAcylation within the multimerization region of galectin 3 plays a role in its secretion. However, surprisingly even the theoretically null mutants where all predicted sites had been deleted or mutated still appeared to retain some level of O-GlcNAcylation (**supplementary Figure s3B**). This would suggest a specific role for O-GlcNAcylation specifically in the multimerization region but it cannot rule out alternate roles in sequestration and release of O-GlcNAcylation other sites of the protein.

In combination with the previous results this would suggest that galectin 3 needs to be first O-GlcNAcylated and then deglycosylated in order to be secreted.

### Galectin 3 secretion is responsive to nutrient conditions via dynamic O-GlcNAcylation

In order to determine if there was a direct link between nutrient conditions and galectin 3 secretion Hela cells were cultured in media with different glucose concentrations (**Figure 8A**). When cultured in low glucose media (1 g/L) and no glucose media (0 g/L) the Hela cells secreted significantly more galectin 3 (∼3 fold and ∼ 10 fold respectively) than cells grown in higher glucose concentrations (2g/L - 4.5 g/L). Western blot analysis showed that O-glcNAcylation increased with glucose concentration while cellular galectin 3 levels were unchanged (**Figure 8B**).

**Figure 8:**
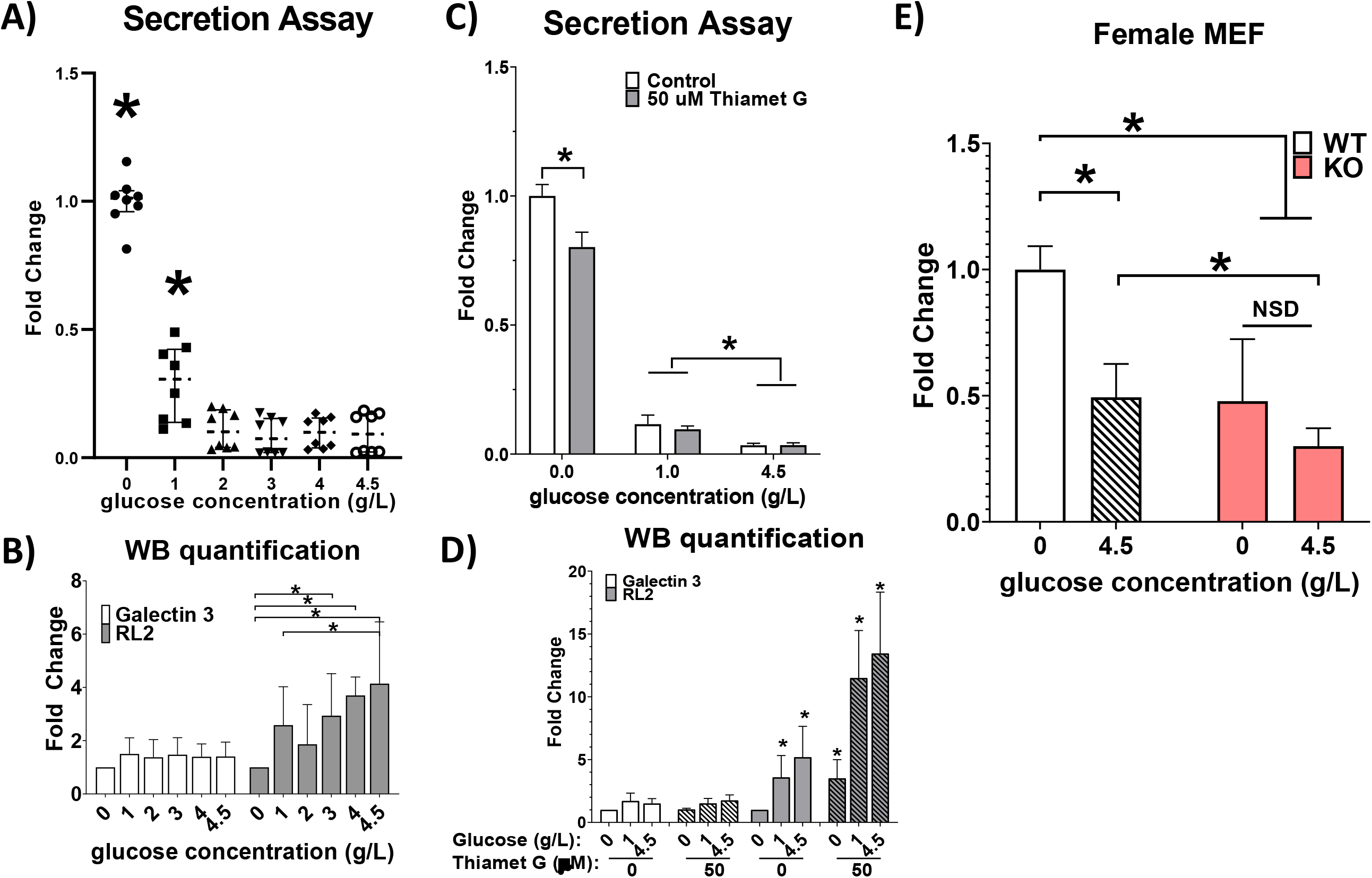
Nutrient conditions have a significant effect on galectin 3 secretion via O-GlcNAcylation. Hela cells grown in different concentrations of glucose were found to secrete significantly more galectin 3 (measured by an ELISA assay) at lower glucose conditions (**A**). LDH release assay performed in parallel shows that there is no significant cytotoxicity at any concentration of glucose except at 0 g/L where there was a small 15% level of LDH release. Quantification of western blots of these Hela cells grown under different glucose conditions shows a glucose dependent increase in O-glcNAcylation corresponding to glucose concentration (**B**). Thiamet G treatment leads to a significant partial ablation of the increased galectin 3 secretion observed at 0 g/L glucose (**C**). Quantification of western blots of these Hela cells grown under different glucose conditions with or without Thiamet G treatment shows a glucose dependent increase in O-glcNAcylation corresponding to glucose concentration and increase in O-GlcNAcylation corresponding to Thiamet G treatment (**D**). Galectin3 secretion was measured by ELISA assay from conditioned media from mouse embryonic fibroblasts from female WT or female OGA KO mice (**E**). The nutrient sensitive increase in galectin3 secretion observed in WT MEFs is absent in OGA KO MEFs. At least three independent experiments were carried out with data expressed as mean ± S.D. (error bars). *, p < 0.05.

The cells were then cultured in media with 0 g/L, 1g/L and 4.5g/L glucose concentrations with or without Thiamet G treatment (**Figure 8C**). When OGA activity was inhibited by Thiamet G treatment there was a significant partial ablation of the nutrient driven increase in galectin 3 secretion in the 0 g/L glucose media condition. This suggests that at least a portion of the nutrient sensitive change in galectin 3 secretion is driven by dynamic O-glcNAcylation. Western blot analysis confirmed that O-glcNAcylation increased with glucose concentration and Thiamet G treatment leads to an increase in O-glcNAcylation levels whereas cellular galectin 3 levels were unaffected (**Figure 8D**).

Thiamet G does not completely inhibit OGA activity. In order to get a clearer picture of the role of dynamic O-glcNAcylation we also looked at Mouse embryonic fibroblasts from wild type and OGA KO mice have a much more dramatic difference between WT and KO lines (**Figure 8D**). In the WT MEF cells we see a significant increase in galectin 3 secretion in cells grown in 0 g/L glucose media (∼2 fold increase) whereas there was no statistically significant change in galectin 3 secretion in KO MEFs grown in 0g/L glucose media. This finding is consistent with a model in which nutrient sensitive changes in galectin 3 secretion are being driven (at least in part) by dynamic O-glcNAcylation.

### O-GlcNAcylation altered galectin secretion can modulate changes in CIE

To connect O-GlcNAcylation regulation of galectin 3 secretion to some of its known extracellular roles we measured the CIE of CD59 after knocking down OGT or OGA (**Figure 9A**). We observed an increase in CD59 internalization when OGT or OGA were knocked down. This increase in CD59 internalization caused by a decrease in galectin 3 secretion is consistent with observed changes in CIE with respect to altered availability of extracellular galectin 3(23). Clathrin mediated endocytosis of transferrin was not affected by these knockdowns again consistent with observed insensitivity of CME to changes in galectin-glycan interactions (**Figure 9B**).

**Figure 9:**
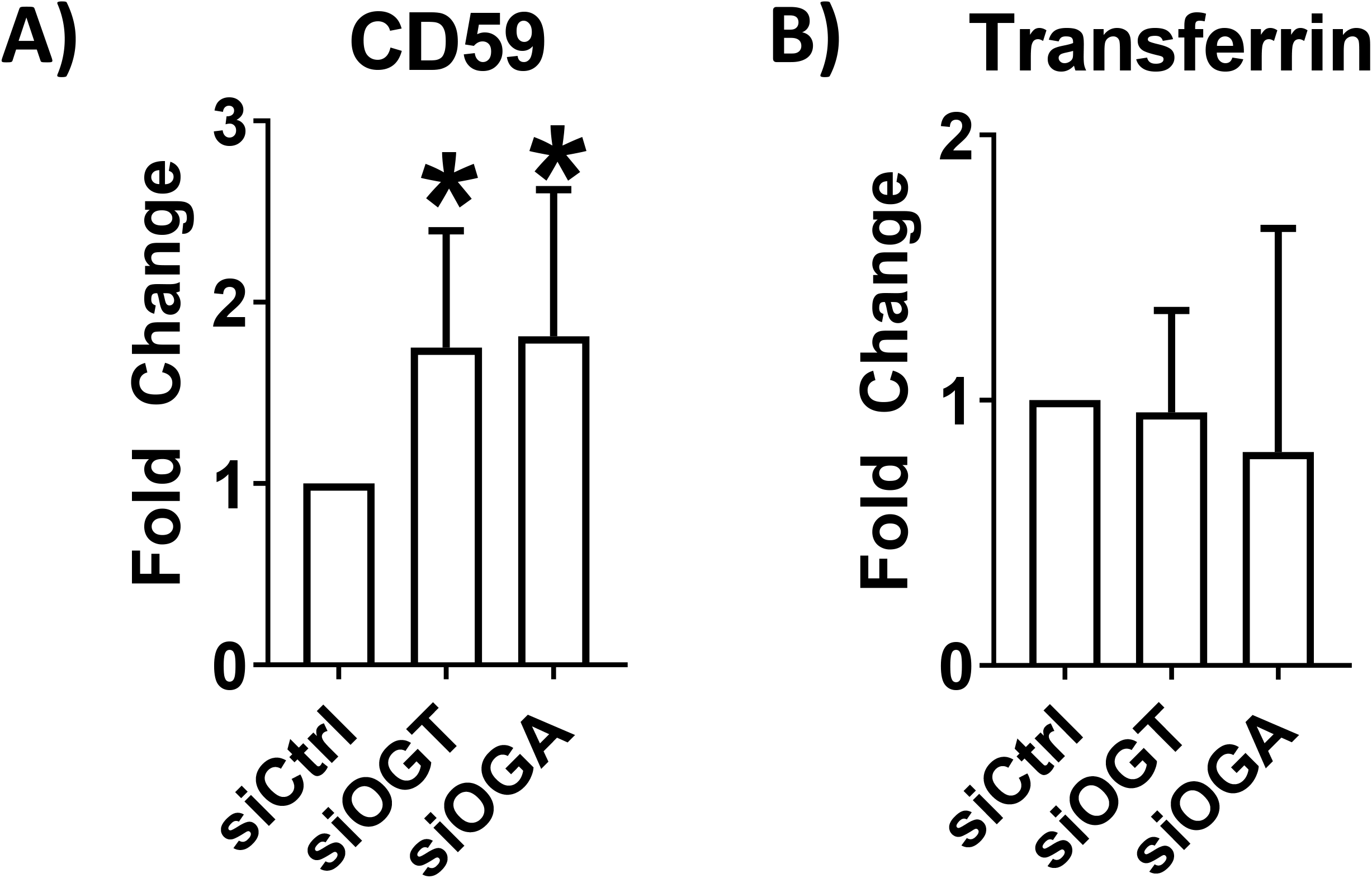
Galectin 3 mediated downstream processes like CIE are altered by disruptions in O-GlcNAc cycling. Clathrin Independent Endocytosis of CD59 measured using and a confocal imaging based antibody internalization assay was found to increase when O-GlcNAc cycling was disrupted by either OGT or OGA knockdown (**A**) whereas clathrin mediated endocytosis of transferrin was not affected (**B**). At least three independent experiments were carried out with representative blots shown and data expressed as mean ± S.D. (error bars). *, p < 0.05.

## DISCUSSION

### Galectin 3 is O-GlcNAcylated

Our data presents the first evidence indicating that a galectin; galectin 3 is *O-*GlcNAcylated (**Figure 2**). This data showing that galectin 3 is O-GlcNAcylated could suggest the possibility that other galectins could also be similarly O-GlcNAcylated. This data could also suggest that other non-canonically secreted proteins like FGF that are synthesized in the cytoplasm could similarly be O-GlcNAcylated.

### Perturbations in O-GlcNAc cycling alter Galectin 3 secretion

Our data shows that secreted galectin 3 is preferentially deglycosylated (**Figure 3**) and that disruption of O-glcNAc cycling leads to altered patterns of galectin 3 secretion. OGA KO MEFs show significant galectin3 secretion defects (**Figure 4**) and siRNA knockdowns (**Figure 5**) and chemical inhibitors (**Figure 6**) are able to alter galectin 3 secretion in Hela cells. This data is the first to our knowledge that shows a link between O-GlcNAcylation and non-classical secretion. Proteins like FGF and other galectins are also secreted by non-classical mechanisms and due to their cytoplasmic synthesis could also have their secretion modulated via O-GlcNAcylation.

Our data also offers the first indication that there is a direct effect of O-glcNAcylation in the unstructured region of galectin 3 on its secretion (**Figure 7**).

While our data indicates that there are potentially indirect effects of altered cellular O-GlcNAcylation and other routes by which altered O-glcNAcylation can modulate galectin 3 secretion (**Figure 2** and **Table 1**) they are beyond the scope of this study. These identified interacting proteins both O-GlcNAcylated and non-glycosylated could be a part of the poorly understood cytoplasmic machinery underlying galectin 3 secretion and would be an interesting focus for future studies.

### Galectin 3 secretion is nutrient sensitive via O-GlcNAcylation

We also provide the first direct evidence that links nutrient sensing to galectin 3 secretion via O-glcNAcylation (**Figure 8**). This is not only important in the context of this study but also in the context of studies that rely on cell culture broadly as we demonstrate that there is a dramatic difference in galectin 3 secretion under high and low glucose media regimes.

Regulation by *O-*GlcNAcylation of galectin 3 secretion provide a molecular mechanism by which cells can integrate numerous signals into feed forward and feedback loops to tune the transduction of nutrient information into functional effects (**Figure 10**). Our data indicates a complex and nuanced role galectin 3 can play as a tunable transducer of nutrient sensing.

**Figure 10:**
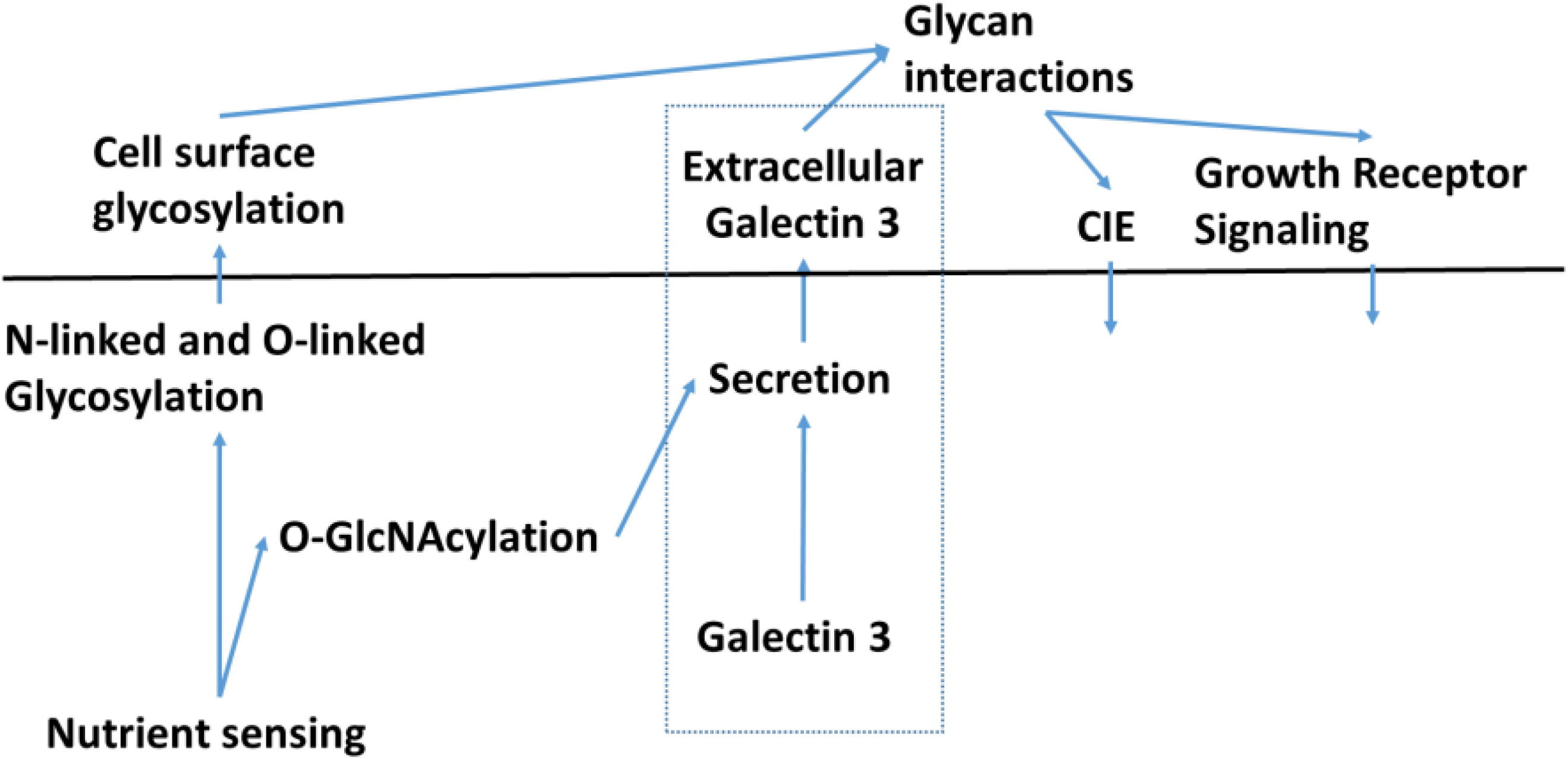
Schematic diagram of the role of galectin 3 as a integrator and tunable transducer of nutrient sensing information. Nutrient sensing information is detected and presented by N-linked glycosylation and O-GlcNAcylation. Galectin 3 is synthesized in the cytoplasm and is O-GlcNAcylated, its O-GlcNAcylation status plays a significant role in its secretion. Extracellular galectin 3 binds to N-linked glycans and drives important cellular effects like CIE and growth receptor signaling. As a result, Galectin 3 can integrate nutrient sensing information from both N-linked glycosylation and O-GlcNAcylation and transduce it into biological effects.

### CIE can be regulated by O-GlcNAc cycling via modulation of galectin 3 secretion

One cellular function that galectin 3 has been shown to play an important role in is CIE(19–23). While CIE is an essential cellular function, it is very poorly understood with little known about how it is regulated(39–44). My previous work has shown that galectin-glycan interactions could be a means by which CIE is modulated(22,23,25). We show that disruption of O-GlcNAcylation by siRNA knockdown of OGT and OGA lead to a decrease in galectin 3 secretion (Figure 5) and that the resulting decrease in extracellular galectin 3 leads to an increase in CD59 CIE (**Figure 9**). This increased internalization of CD59 was consistent with the trends we had previously observed when galectin 3 was knocked down in Hela cells(23). By uncovering the role of O-GlcNAcylation and nutrient sensing in regulating galectin 3 secretion and by directly linking O-GlcNAcylation disruption to modulation of CIE of CD59 we offer the first indication that CIE could be nutrient sensitive.

### Dysregulation of galectin 3 in disease

In addition to its unique biological niche, galectin 3 dysregulation has also been observed in a wide variety of disease contexts from cancer to diabetes and heart disease(45–52). Hence, understanding the molecular mechanisms driven by galectin-glycan interactions and how galectin 3 secretion is modulated will provide us with valuable insights into how galectins and glycosylation drive the progression of several diseases.

## EXPERIMENTAL PROCEDURES

### Cell culture, reagents, and antibodies

HeLa and Mouse Embryonic Fibroblast (MEF) cells were grown in Dulbecco’s modified Eagle’s medium (Lonza) supplemented with 10% heat-inactivated fetal bovine serum (Atlanta Biologicals), 1.0% 100× glutamine solution (Lonza), and 1.0% 100× penicillin/streptomycin antibiotic solution (Lonza). Cells were maintained at 37 °C in a humidified, 5% CO_2_-containing atmosphere.

For GlcNAc treatment, a 100 mM stock solution of GlcNAc (Sigma-Aldrich) was prepared in sterile complete culture medium, cells were then seeded on coverslips or on tissue culture plastic in 60-mm culture plates in 5.0 ml of culture medium at a density of 3 × 10^5^ cells/plate, and the appropriate volume of GlcNAc stock solution was added to each well to achieve the desired sugar concentrations. Cells were typically incubated for 48 h with the sugar. Similarly, for glucose treatment controls, a 100 mM stock solution of glucose (J.T. Baker) in sterile complete culture medium was prepared and used.

For Thiamet G treatments, a 2 μg/ml solution of Thiamet G (Sigma-Aldrich) in sterile DMSO was prepared, and the cells were incubated in the Thiamet G solution for 48-72 h before the start of the experiment.

For OSMI-1 treatments, a 2 μg/ml solution of OSMI-1 (Sigma-Aldrich) in sterile DMSO was prepared, and the cells were incubated in the OSM-1 solution for 48-72 h before the start of the experiment.

Monoclonal antibodies directed toward CD59 (clone p282/H19) was from Biolegend. A monoclonal antibody directed toward galectin 3(clone A3A12) was from ThermoFisher. A polyclonal antibody that detects galectin 3 (ab53082) was from Abcam. A polyclonal antibody that detects Ezrin (catalog no. 3145S) was from Cell Signaling. A monoclonal antibody that detects serine or threonine linked O-Linked N-Acetylglucosamine-RL-2 (ab2739) was from abcam. Anti-OGT (ab177941), anti-OGA(ab124807) and anti-Tubulin (ab6046) were also from abcam. An agarose conjugated mouse monoclonal anti-galectin 3 (sc-32790 AC) was from santa cruz biotechnology. Alexa 594–conjugated transferrin and all secondary antibodies conjugated to Alexa Fluor 594, 488, 647, 680, and 800 were purchased from Molecular Probes.

### Antibody internalization assay

After the indicated incubations, transfections, and pretreatments, cells on coverslips were placed face up on parafilm, and 50 μl of conditioned complete medium containing primary antibody at concentrations of 5 μg/ml for CD59 (Biolegend) or 10 μg/ml Alexa Fluor 488–linked transferrin (Molecular Probes) was added. Coverslips were then incubated either on ice or at 37 °C for 30 min. 10-min incubations were used for transferrin-labeled coverslips. After this, one set of coverslips that was incubated at 37 °C was fixed in 2% formaldehyde (representing total antibody bound both internally and to the surface, *T*_tot_), and the other set of coverslips that was held at 37 °C (representing antibody internalized, *T*_int_) as well as the set that was kept on ice (null control, *T*_0_) were acid-washed for 40 s using a solution of 0.5% acetic acid, 0.5 M NaCl, pH 3.0, after which these sets of coverslips were fixed in 2% formaldehyde. The cells were blocked for 30 min using a blocking solution of 10% FBS, 0.02% sodium azide in PBS. The coverslips were then labeled at room temperature for 1 h with Alexa Fluor 488–conjugated goat anti-mouse secondary antibody (Molecular Probes), 0.2% saponin, and 2 μg/ml HCS cell mask deep red stain (Molecular Probes) in blocking solution. The coverslips were then rinsed three times in blocking solution and a final time with PBS, and they were then mounted on glass slides.

Coverslips were imaged at room temperature using a confocal microscope (LSM 780 FCS, Carl Zeiss) with a 40× PlanApo oil immersion objective and 488- and 633-nm laser excitation. For each coverslip, six positions were imaged with 2 × 2 tiling with the pinhole kept completely open. For each condition across the six tiled images, at least 100 cells were imaged. All images for each antibody were taken with identical acquisition parameters, which are set based on the control’s *T*_tot_ coverslip and tuned such that the signal was within the dynamic range.

The MetaMorph application (Molecular Devices) was used to quantify the percentage of antibody internalized. Confocal images were separated into the two different channel colors. An antibody channel threshold was set for each condition using its *T*_0_ coverslip images. The *T*_int_ and *T*_tot_ conditions were then thresholded, and the integrated signal intensity was measured for the antibody channel. The cell mask channel was autothresholded, and the threshold area was measured. The integrated signal intensity for the antibody was then normalized to the threshold area for each image. The percentage internalized is then calculated with the equation, % internal = 100 ×*T*_tot/area_/*T*_int/area_. Finally, the internal percentages were normalized to the untreated control.

### Galectin 3 ELISA secretion assay

Cells were seeded on a 6-well plate at a density of 3 × 10^5^ cells/well in 1 ml of complete medium in quadruplicate with relevant siRNA transfections and chemical inhibitor treatments. After a 48 h or 72h incubation, the supernatants (*i.e.* conditioned medium) were collected and spun down at 3000 × *g* for 10 min to remove any cell debris. Supernatants were then analyzed for galectin 3 content using a human galectin 3 ELISA kit (Abcam, ab 188394) as recommended by the manufacturer. Cells in the well were trypsinized to detach and counted. Galectin 3 in the supernatant was normalized to cell number.

### siRNA transfection

Cells were seeded in antibiotic-free complete medium and transfected with a 50 nM final concentration of OGT (Dharmacon, ON-TARGET plus human OGT (8473) siRNA-smartpool), OGA (Dharmacon, ON-TARGET plus human MGEA5 (10724) siRNA-smartpool) or nontargeting siRNA (Dharmacon, ON-TARGETplus nontargeting pool) using Lipofectamine RNAiMAX (Invitrogen) as recommended by the manufacturer. Cells were used in experiments 48-72 h after transfection.

### Western blot analysis

Proteins obtained from cells were analyzed by Western blotting. Briefly, after the appropriate treatment (transfection, siRNA treatment, GlcNAc incubation, etc.), cells were washed three times in cold PBS, harvested by using a tissue culture cell scraper, and pelleted by centrifugation at 4 °C (1200 × *g* for 10 min). Pellets were solubilized in RIPA buffer (Abcam) protein levels were measured and normalized using a BCA assay (Thermofisher). Proteins were separated by 10–20% SDS-PAGE, transferred to nitrocellulose paper, and probed with the following commercial antibodies: anti RL-2 (Abcam), anti-tubulin (Abcam), anti-OGT (Abcam), anti-OGA(Abcam), anti-Ezrin (Cell Signaling) and anti-galectin 3 (ThermoFisher). Membranes were imaged using LI-COR Odyssey IR imager. Protein bands were quantified using the Image Studio Lite software.

### Galectin 3 immunoprecipitation assay

After a 48-h incubation with or without GlcNAc treatment, cells were rinsed with ice-cold PBS and lysed in 1 ml of ice-cold RIPA buffer (150 mM NaCl, 1% Nonidet P-40, 0.5% sodium deoxycholate, 0.1% SDS, 50 mM Tris, pH 8.0) with protease inhibitor mixture (Sigma-Aldrich). Lysates were centrifuged at 13,300 rpm for 10 min. Supernatants were transferred to fresh Eppendorf tubes. The samples were immunoprecipitated using 15 μg/ml agarose conjugated anti-galectin 3 (Santa cruz biotechnology) or agarose Protein G beads (GE Healthcare) only. The beads were preblocked for 1 h on ice with 3% BSA (Sigma-Aldrich) in RIPA buffer. After immunoprecipitation, beads were rinsed three times with RIPA buffer and boiled for 10 min in 1× sample-loading buffer, and proteins were separated by 10–20% SDS-PAGE, transferred to nitrocellulose paper, and probed with the following antibodies: anti-Ezrin (Cell Signaling), anti-galectin 3 (Abcam), and anti-RL-2 (Abcam). Membranes were imaged using a LI-COR Odyssey IR imager.

### Plasmids, primers and transient transfection

GFP-tagged galectin 3 (pEGFP-hGal3 (Addgene, plasmid no. 73080) was mutated using the Q5 site directed mutagenesis kit (New England Biolabs, catalog number E0554S) and designed primers according to the manufacturer’s instructions. Required mutations, deletions, insertions and combinations of these were generated using the following primers: *GFP-Gal 3 Del (84S-104T deletion)* Forward primer: 5’-GGCCCCTATGGCGCCCCT-3’, Reverse primer: 5’- GGGTGGCCCTGGGTAGACTC-3’. *GFP-Gal 3 1 mutation (243 T/A)* Forward primer: 5’- CATAGACCTCgccAGTGCTTCAT -3’, Reverse primer: 5’- TCACCAGAAATTCCCAGTTTG -3’. *GFP-Gal 3 Deletion null(84S-104T deletion and 243 T/A):* using the confirmed GFP-Gal 3 Del mutant the primers for GFP-Gal 3 1 mutation (243 T/A) were used. *Insertion (84S-104T, 84S/A,91S/A,92S/A,104T/A)* Forward primer: 5’- gccagccgcccccggagcctaccctgccactGGCCCCTATGGCGCCCCT -3’, Reverse primer: 5’-tgtccggcggctgggtaggccccagggccggcGGGTGGCCCTGGGTAGACTC-3’. *GFP-Gal 3 4 Mutation (84S/A,91S/A,92S/A,104T/A)* using the confirmed GFP-Gal 3 Del mutant the primers for Insertion were used. *GFP-Gal 3 Mutation null (84S/A,91S/A,92S/A,104T/A and 243 T/A)* using the confirmed GFP-Gal 3 Deletion null mutant the primers for Insertion were used. *Control silent (124G/G and 125G/G)* Forward primer: 5’- GCCTTTGCCTggagggGTGGTGCCTC -3’, Reverse primer: 5’- AGGTTATAAGGCA CAATCAGTGGCCC -3’. *Control random subsititution (124G/A)* Forward primer: 5’- GCCTTT GCCTgcgGGAGTGGTGC -3’, Reverse primer: 5’- AGGTTATAAGGCACAATCAGTGGC -3’ *Sequencing Primers* Forward primer: 5’- CGGACTCAGATCTATGGCAGAC -3’, Reverse primer: 5’- AAACCTCTACAAATGTGGTATGGC -3’.

Cells were seeded in complete medium. After a 24-h incubation, cells were transfected with GFP-tagged galectin 3 (pEGFP-hGal3) or mutants of it, or pEGFP-N3 (Clontech, catalogue no. 6080-1) at 2.5 μg of DNA/60-mm well using Xtremegene9 (Roche Diagnostics) as recommended by the manufacturer. Experiments were performed 48-72 h after transfection.

### In-vitro OGT assay

The reaction mixture was prepared by mixing 2 μg of recombinant OGT, 3 μg of recombinant galectin 3(Abcam, ab89487), 1 μL of 2M UDP-GlcNAz and making the volume up to 50 μL with OGT assay buffer (50 mM Tris–HCl, pH 7.5, 1 mM dithiothreitol, 12.5 mM MgCl_2_). Negative controls were prepared by removing either OGT, galectin 3 or UDP-GlcNAz from the reaction mixture. A positive control was prepared by substituting casein kinase II(NEB, P6010S) for galectin 3 in the reaction mixture.

The OGT assay mixtures were incubated for ∼90 minutes in a 37°C water bath. The reaction mixtures were then added to 10K filters and spun at 14000 rpm for 30 min. The filters were washed with 100μl of PBS and spun down at 14000xg for 10 min twice. To recover the protein the membranes were flipped and spun down at 1000xg for 3 minutes. 1 μl of biotin phosphine was added and incubated for 1 hour at 40°C. The samples were then boiled for 10 min in 1× sample-loading buffer, and proteins were separated by 10–20% SDS-PAGE, transferred to nitrocellulose paper, blocked in odyssey blocking buffer for 1 hour and probed with streptavidin IR800 and anti-galectin 3 (Abcam). Membranes were imaged using a LI-COR Odyssey IR imager.

### In-silico O-glcNAcylation site prediction

The galectin 3 sequence was used on the YinOYang 1.2 server to predict site of O-GlcNAclation (http://www.cbs.dtu.dk/services/YinOYang/)(53, 54). The server generates a list of predicted sites with varying degrees of confidence.

### Mass spectrometric analysis

After Galectin 3 immunoprecipitation the samples were resolved on SDS-NuPAGE gel and stained with Simply Safe Blue (Invitrogen). Multiple protein bands were excised, destained with 50% methanol in 50mM bicarbonate, reduced with 20 mM DTT at 57 °C for 45 min and alkylated with 50 mM iodoacetamide in the dark at room temperature for 30 min, and then were digested with 5ng/ul of trypsin (Promega) at room temperature overnight according to the manufacturer’s protocol. The resulting peptides were dried and reconstituted in 0.1% formic acid and desalted by C18 ziptip (EMD Millipore) according to the manufacturer’s protocol. The eluted peptides were dried again and dissolved with sample loading buffer (0.1% formic acid, and 5% AcN) for LC-MS/MS analysis.

The digested peptides were analyzed by reverse phase LC-MS/MS on an Orbitrap Fusion Lumos mass spectrometer (Thermo Scientific) coupled to a Dionex UltiMate 3000 -nLC (Thermo Scientific) liquid chromatography system. The instrument was operated in data-dependent mode. The data were searched against a UniProt human database (downloaded 01/2018), using SEQUEST within the Proteome Discoverer 1.4.

### GFP-Trap Immunoprecipitation of GFP tagged galectin 3 mutants

After a 48-h transfection with GFP-tagged galectin 3 mutants cells were rinsed with ice-cold PBS and lysed in 1 ml of ice-cold RIPA buffer (150 mM NaCl, 1% Nonidet P-40, 0.5% sodium deoxycholate, 0.1% SDS, 50 mM Tris, pH 8.0) with protease inhibitor mixture (Sigma-Aldrich). The conditioned media was also collected in some experiments and spun down at 3000 × *g* for 10 min to remove any cell debris, supernatants were transferred to fresh eppendorf tubes. Cell lysates were centrifuged at 13,300 rpm for 10 min. Supernatants were transferred to fresh eppendorf tubes. The samples were immunoprecipitated using 25 μL bead slurry of GFP-Trap Agarose (Chromotek). The beads were preblocked for 1 h on ice with 3% BSA (Sigma-Aldrich) in RIPA buffer. After immunoprecipitation, beads were rinsed three times with wash buffer (10 mM Tris/Cl pH 7.5, 150 mM NaCl, 0.05 % Nonidet™ P40 Substitute, 0.5 mM EDTA) and boiled for 10 min in 1× sample-loading buffer, and proteins were separated by 10–20% SDS-PAGE, transferred to nitrocellulose paper, and probed with the following antibodies: anti-Ezrin (Cell Signaling), anti-galectin 3 (Abcam), and anti-RL-2 (Abcam). Membranes were imaged using a LI-COR Odyssey IR imager.

### GFP-tagged galectin 3 secretion assay

Cells were seeded on a 12-well plate at a density of 5 × 10^4^ cells/well in 1 ml of complete medium in quadruplicate. Cells were transfected with GFP-tagged galectin 3 (Addgene) or a mutant variant of it. After 24 h, the medium was replaced with 500 μl of FluoroBrite Dulbecco’s modified Eagle’s medium (Life Technologies) supplemented with 10% heat-inactivated fetal bovine serum (Atlanta Biologicals), 1.0% 100× glutamine solution (Lonza), and 1.0% 100× penicillin/streptomycin antibiotic solution (Lonza). 72 h post-transfection, the supernatants were collected and spun down at 3000 × *g* for 10 min to remove any cell debris, and 200 μl of the supernatant was transferred to a black 96-well plate in duplicate. The cells were also detached by incubating with 0.25% trypsin for 1 min at 37 °C. Cells were collected, spun down, and resuspended in 500 μl of PBS. 200 μl of the cell suspensions were also transferred to a black 96-well plate in duplicate. Bulk fluorescence in each well was measured using a Synergy H1 multimode microplate reader (BioTek). After background-subtracting any fluorescence from the medium, PBS, and cell autofluorescence, the fluorescence from the supernatant wells was normalized to the fluorescence in the cell suspension wells.

### Statistical analysis

At least three independent biological replicates (independent experiments) were carried out for each experiment, and data were expressed as mean ± S.D. In experiments with multiple comparisons, statistical significance was determined using a one-way ANOVA with a Dunnett’s post-test to compare means of different samples with the control or a Bonferroni post-test to compare specific pairs of columns. In experiments with only two conditions, an unpaired Student’s *t* test was used to determine statistical significance. The null hypothesis was rejected in cases where *p* values were <0.05.

## SUPPORTING INFORMATION

This article contains supporting information.

## ACKNOWLEDGEMENTS

The mass spectrometry experiments were carried out by Yanling Yang and Marjan Gucek at the Proteomics Core Facility, NHLBI, NIH, Bethesda, Maryland, USA. Recombinant OGT was expressed and purified by Agata Steenackers.

## FUNDING AND ADDITIONAL INFORMATION

This work was supported by NHLBI, National Institutes of Health, intramural research program grant Hl006130 and by NIDDK, National Institutes of Health, intramural research program grant ZIADK060103.

